# Hero11 Unlocks TDP-43 Condensate Fluidity via Targeting Inter-Helical Interactions

**DOI:** 10.64898/2026.05.11.724201

**Authors:** Cheng Tan, Jaewoon Jung, Yuji Sugita

## Abstract

Biomolecular condensates formed by intrinsically disordered proteins (IDPs) rely on a balance of sequence-encoded interactions and secondary-structure elements. TDP-43, a disease-associated protein, undergoes liquid–liquid phase separation (LLPS) through its low-complexity domain, whereas Hero11 has been proposed to modulate its condensate properties. However, the molecular mechanisms by which Hero11 affects the internal organization and dynamics of TDP-43 condensates remain unknown. Here, using multi-microsecond explicit-solvent all-atom simulations spanning single chains to =~100-chain condensates, we show that the TDP-43 α-helix, which is only marginally stable in isolation, becomes a major structural hub within the condensate, forming a percolated helix–helix interaction network whose contact lifetimes are substantially longer than those of the surrounding disordered contacts. Hero11 selectively dismantles this network: it binds preferentially near the helical region, reduces the helix–helix coordination number, and shortens helix–helix contact lifetimes. This targeted disruption lowers condensate density, increases both water and ion infiltration, and enhances TDP-43 diffusion within the dense phase. Notably, dimer simulations reveal that the interactions between TDP-43 and Hero11 are too weak to persist under dilute conditions, indicating that the regulatory effect emerges only through multivalent contacts in the condensed phase. These results establish the α-helix as a selectively vulnerable structural element within the TDP-43 condensate and provide an atomic-level mechanism for how a highly charged disordered protein can tune condensate material properties by targeting its longest-lived interaction nodes.

## Introduction

Biomolecular condensates formed via liquid–liquid phase separation (LLPS) organize cellular biochemistry by compartmentalizing proteins and nucleic acids without the use of surrounding membranes^1,2^. A defining feature of these condensates is their dynamic, liquid-like character, which allows rapid exchange of components with the surrounding environment. However, under pathological conditions, condensates can mature into solid-like aggregates that are associated with a range of neurodegenerative diseases^3^. TDP-43, an RNA-binding protein whose C-terminal low-complexity domain (LCD) is the primary driver of phase separation^4^, is a prominent example: its condensates are functional under normal conditions but are prone to forming irreversible amyloid fibrils linked to neurodegeneration^5,6^. Understanding the molecular forces that maintain the liquid-like state and prevent pathological solidification is therefore a central question in the biophysics of biomolecular condensates.

The phase separation of TDP-43 LCD is driven by a combination of interaction modes typical of intrinsically disordered regions, including aromatic and hydrophobic contacts, electrostatic interactions, and hydrogen bonding. Among these, a conserved α-helical region (residues =~320–340 in the full-length protein) has attracted particular attention^4,7,8^. NMR studies have shown that this region is transiently helical in the monomeric state but becomes stabilized and extended upon self-association, and that the resulting intermolecular helix–helix contacts contribute significantly to phase separation^4^. Subsequent work demonstrated that designed mutations enhancing helical propensity at conserved glycine positions markedly promote LLPS, while ALS-associated mutations in the same region disrupt helix–helix interactions and reduce phase separation^7^. More recently, structural characterization has provided direct evidence for helix-mediated multimerization of the TDP-43 C-terminal domain^8^. These findings establish the α-helical region as a key interaction element within the otherwise disordered LCD, but how this helical element functions within the dense, crowded environment of a condensate, or in the presence of regulatory molecules, has not been characterized at the atomic level.

The physical state of protein condensates can be modulated by interacting molecules. RNA, for example, has been shown to buffer the phase separation of prion-like RNA-binding proteins at high concentrations and to antagonize pathological phase transitions of TDP-43^9,10^. More recently, a class of intrinsically disordered proteins termed Hero (heat-resistant obscure) proteins has been identified as a distinct type of condensate regulator^11^. Hero proteins are characterized by high charge density and disordered conformations, and function without ATP. Hero11, a small and highly positively charged member of this family, suppresses TDP-43 aggregation in cellular models^11^ and promotes extended conformations of TDP-43 LCD as observed by single-molecule FRET^12^. However, the molecular mechanism by which Hero11 modulates TDP-43 condensates remains unknown. In particular, it is unclear whether Hero11 acts through nonspecific electrostatic effects, as might be expected from a highly charged polymer, or whether it engages specific structural features of the condensate, such as the helical interaction network described above.

Characterizing these interactions at the atomic level is challenging for most experimental techniques, and computational approaches have therefore played an important role. Coarse-grained (CG) simulations, in particular, have been widely used to capture the phase diagrams and mesoscale organization of IDP condensates^13,14^. In our previous study, we employed CG simulations of TDP-43 LCD and Hero11 mixtures and identified electrostatic repulsion among Hero11 chains as a key factor in destabilizing the TDP-43 condensate, leading to reduced interchain contacts and accelerated protein diffusion^15^. However, CG models inherently lack the resolution to describe several features that may be central to the mechanism of condensate regulation: transient secondary structure formation and its environment-dependent conformational changes, specific side-chain geometries at interaction interfaces, and the behavior of explicit water molecules and ions within the condensed phase. Consequently, it remains unknown how Hero11 affects the internal structural organization of the TDP-43 condensate at the atomic level, and whether the helical interaction network identified by experiment plays a role in this process.

Here, we address these questions by performing large-scale, all-atom molecular dynamics simulations of TDP-43 LCD condensates in the presence and absence of Hero11, accumulating over 20 μs of trajectory data. We find that within the homotypic condensate, the α-helical region of TDP-43 is stabilized by the crowded environment and forms a percolated helix–helix interaction network that serves as a structural hub of the dense phase. Hero11 preferentially binds near this helical region and is associated with reduced helix– helix coordination, decreased helical propensity, increased water and ion infiltration, and enhanced protein diffusion. In contrast, dimer simulations in dilute solution show that the pairwise TDP-43–Hero11 interaction is weak and transient, indicating that the regulatory effect of Hero11 depends on the multivalent environment of the condensed phase. Together, these results provide an atomic-level picture of how a highly charged disordered protein modulates the internal organization and physical properties of a biomolecular condensate.

## Results

### Simulation Systems

We initialized our all-atom (AA) simulations using equilibrated configurations derived from our previous coarse-grained (CG) simulations based on the HPS model^16,17^. Specifically, we extracted representative snapshots from the center of a stable pure TDP-43 system containing 100 chains, as well as mixed condensates consisting of 100 TDP-43 chains and a varying number of Hero11 chains (*n* = 17, 18, 19 for the three independent replicas).

The CG beads were backmapped to all-atom coordinates using the PULCHRA software package^18^ to reconstruct backbone and side-chain geometries. These reconstructed structures were then organized into three distinct simulation setups (Figure 1): (i) Single-chain systems, representing the infinite dilution limit, where individual TDP-43 and Hero11 chains were solvated in cubic boxes (edge length of 220Å for TDP43 and 180Å for Hero11) to characterize their intrinsic conformational preferences; (ii) Dimer systems, designed to explicitly probe possible specific intermolecular contacts in cubic boxes with edge length of 235Å; and (iii) Condensate systems, where full collections of reconstructed chains were placed in elongated boxes (180 × 180 × 600 Å^3^) to accommodate the slab geometry for capturing bulk properties.

**Figure 1.**
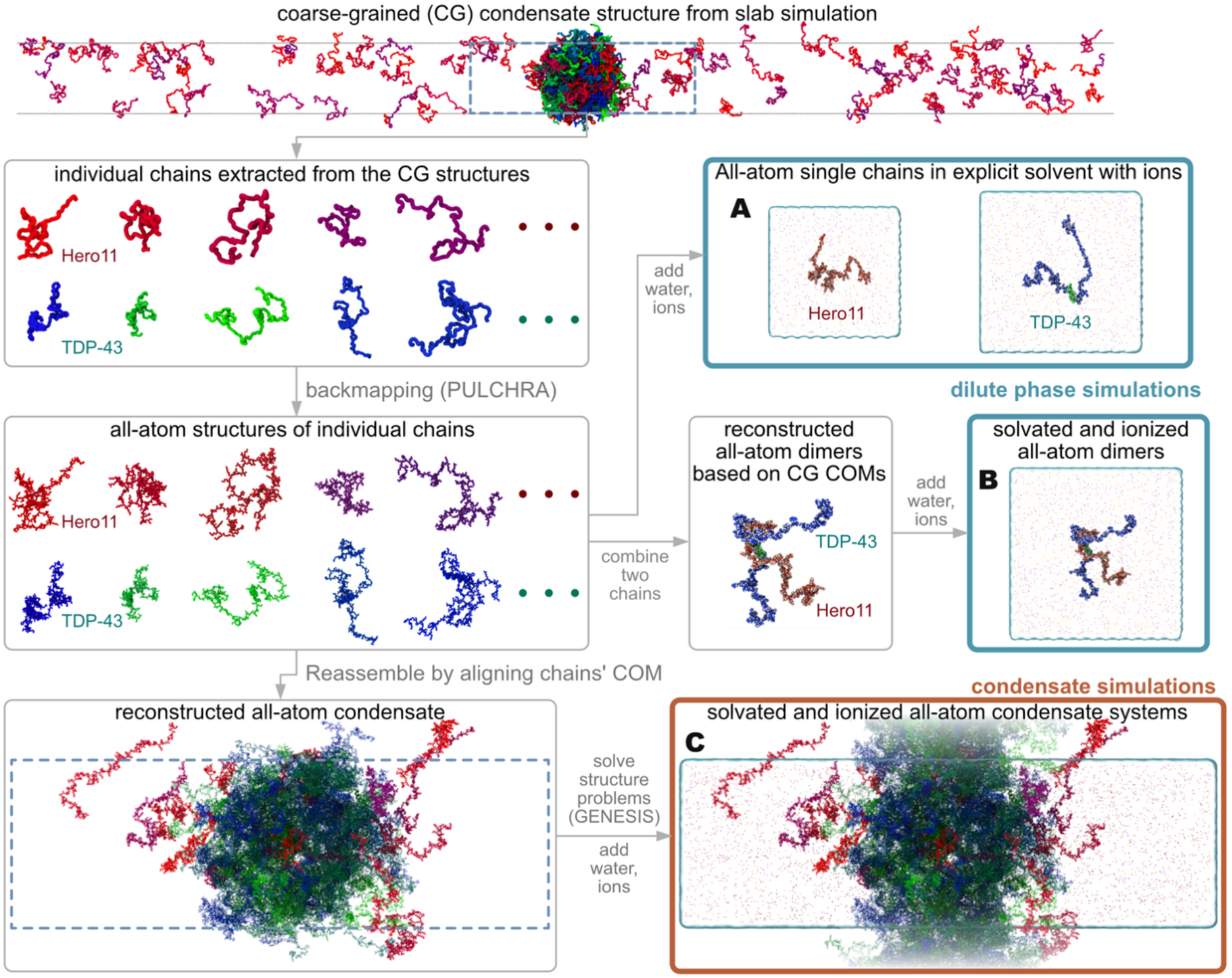
Workflow for generating all-atom (AA) initial configurations from coarse-grained (CG) simulation data. The central region of the CG condensate was used to extract coordinates for individual Hero11 and TDP-43 chains, which were then backmapped to AA resolution using PULCHRA. From this pool of structures, we constructed: (A) single-chain systems in explicit solvent; (B) heterodimer systems in explicit solvent identified from proximal pairs in the CG trajectory; and (C) full condensate systems (homotypic TDP-43 and heterotypic TDP-43/Hero11) reassembled by restoring the original CG center-of-mass (COM) coordinates. Structural clashes were resolved using GENESIS before solvation and ionization.

All systems were solvated with explicit water, and ions were added to neutralize the charge and reach a physiological salt concentration of 150 mM. To address structural artifacts inherent to the backmapping process, specifically ring penetrations, steric clashes, and chirality errors, we employed the energy minimization and structural refinement protocols available in GENESIS to ensure stereochemical validity^19^. These setups enable a systematic comparison of TDP-43 properties and Hero11 regulations across different density regimes, from isolated chains to the dense liquid network.

### Conformational Properties of Single Chains

#### TDP-43 Adopts Compact Conformations with Transient Helicity

In the single-chain simulations, TDP-43 adopts a relatively compact conformation. Even when initialized from an extended state, its radius of gyration (*R*_*g*_) rapidly decreases and fluctuates around =~27 Å (Figure 2A), indicating an intrinsic tendency toward chain compaction, as illustrated in the representative snapshots (Figure 2C). The intramolecular contact probability map (Figure S2A) shows that contacts are predominantly short-range, distributed along the sequence diagonal. However, a small number of long-range contacts persist over hundreds of nanoseconds (Figure S2B, S2C). The presence of persistent long-range contacts is consistent with, and likely contributes to, the compact conformation adopted by the isolated chain.

**Figure 2.**
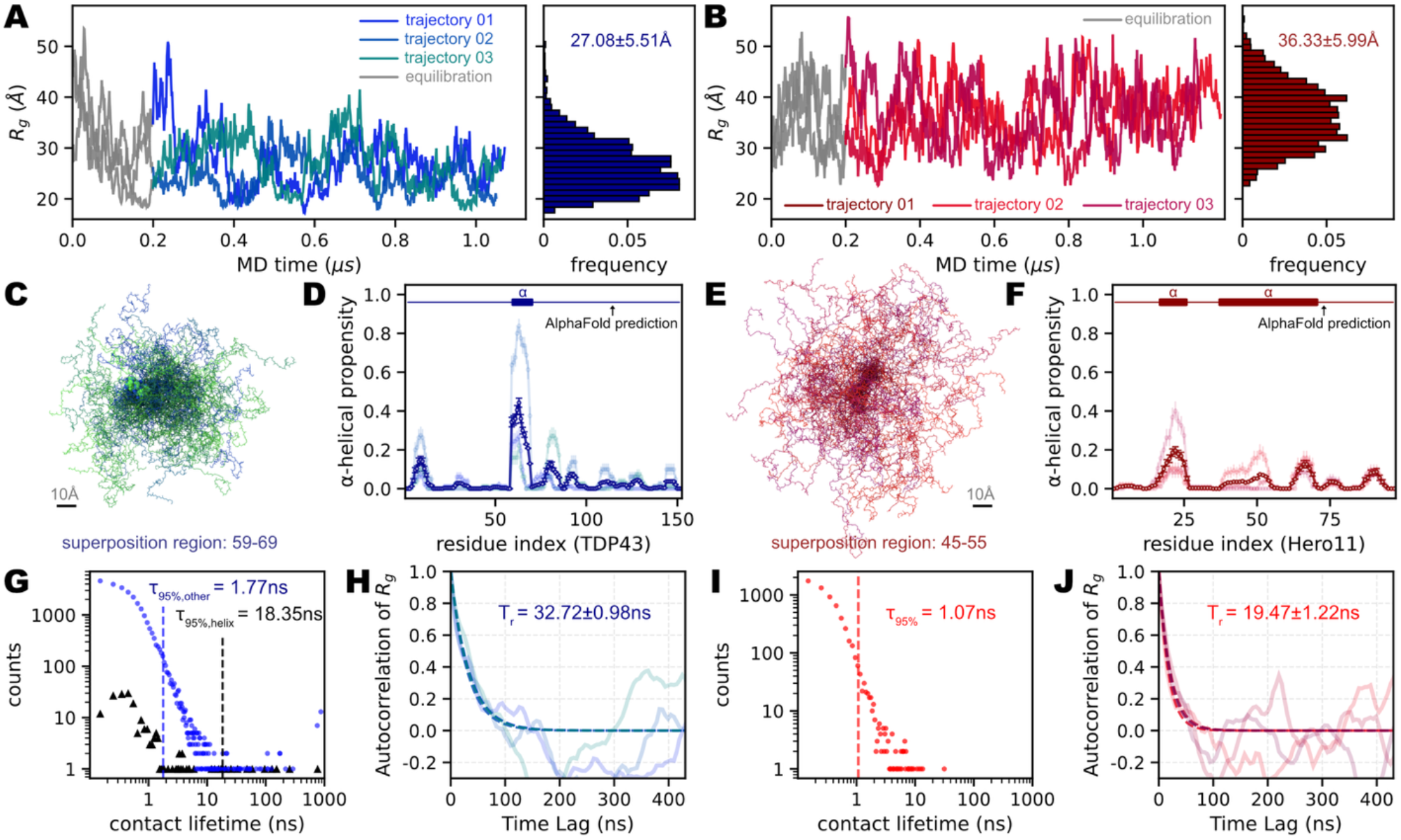
Conformational ensembles and dynamics of single-chain TDP-43 and Hero11. (A, B) Time series and probability distributions of the radius of gyration (*R*_*g*_) for TDP-43 (A) and Hero11 (B). Three independent =~1.2 µs replicas are overlaid. Gray regions indicate the initial 200 ns equilibration. Mean values indicate a compact state for TDP-43 (=~27 Å) and an extended state for Hero11 (=~36 Å). (C, E) Superimposed snapshots from production trajectories, illustrating the collapsed conformation of TDP-43 (C) versus the expanded conformation of Hero11 (E). (D, F) Residue-wise α-helical propensity. TDP-43 forms a transient helix at residues 59–69 (D), whereas Hero11 shows negligible secondary structure (F). Semi-transparent lines represent individual replica trajectories; the solid line with error bars indicates the mean ± s.e.m. across replicas. AlphaFold-predicted helical regions are indicated above each panel. (G, I) Distributions of intramolecular contact lifetimes. For TDP-43 (G), black triangles denote contacts within the helical region and blue circles denote other non-local contacts; vertical dashed lines indicate the 95th-percentile contact lifetime (*τ*_95%_) for each contact class. For Hero11 (I), all contacts decay rapidly, with a single *τ*_95%_ of 1.07 ns. (H, J) Time-autocorrelation functions of Rg quantifying structural relaxation. Semi-transparent lines show individual replica ACFs; dashed lines indicate single-exponential fits to each replica. Reported Tr values represent the mean ± s.d. across replicas.

The α-helical region spanning residues 60–71 exhibits a moderate helical propensity (=~50%, Figure 2D) but is intrinsically unstable in dilute solution, undergoing repeated folding and unfolding events across all three independent trajectories (Figure S2D). The contact lifetime analysis (Figure 2G) shows that interactions within this helical region have longer lifetimes than the non-specific contacts in the surrounding disordered regions, as reflected by the higher 95th percentile values. This contrast between the transient nature of helix formation and the relatively long-lived helix-internal contacts suggests that once formed, the helix provides a locally cohesive structural element, but it is not maintained stably in the absence of intermolecular interactions. The autocorrelation function of *R*_*g*_ decays with a relaxation time of =~33 ns (Figure 2H), indicating that conformational fluctuations persist over tens of nanoseconds.

#### Hero11 Behaves as a Highly Expanded Polyelectrolyte

Hero11 adopts an extended conformation in solution. Its *R*_*g*_ distribution is centered at higher values (=~36 Å) with broader fluctuations than TDP-43, despite the shorter chain length (Figure 2B, E). This behavior is consistent with the high net positive charge of Hero11, which generates strong intrachain electrostatic repulsion and favors chain expansion. Although AlphaFold predicts two α-helical regions in Hero11, including a long helix spanning residues =~38–72 with high confidence (pLDDT > 90; Figure S1B), our microsecond explicit-solvent simulations show that Hero11 samples considerably more disordered conformations than TDP43 over the same timescale. (Figure 2F). This discrepancy is expected for a protein with such high charge density, as the electrostatic self-repulsion that extends the chain also disfavors the formation of compact helical segments. The intramolecular contacts of Hero11 are transient (Figure 2I), and the rapid decay of the contact autocorrelation function (Figure 2J) indicates fast internal relaxation dynamics. These single-chain properties of TDP-43 and Hero11, namely, compact and cohesive versus extended and self-repulsive, set the stage for understanding their distinct roles in the condensate environment examined below.

### TDP-43−Hero11 affinity: too weak to sustain a dimer in dilute phase

To examine the pairwise interaction between TDP-43 and Hero11, we performed all-atom MD simulations of TDP-43–Hero11 heterodimers in explicit solvent. Four independent trajectories, each exceeding 1 μs, were initiated from configurations in which TDP-43 and Hero11 were placed in close proximity (Figure 1B). Over the course of the simulations, the center-of-mass distance (*d*_*COM*_) between the two chains increased, and the number of intermolecular contacts (*n*_*cnt*_) decreased in most trajectories, indicating that the heterodimer tends to dissociate under dilute conditions (Figure 3A). This result suggests that the pairwise interaction between TDP-43 and Hero11 is intrinsically weak in the absence of the multivalent environment present in the condensed phase.

**Figure 3.**
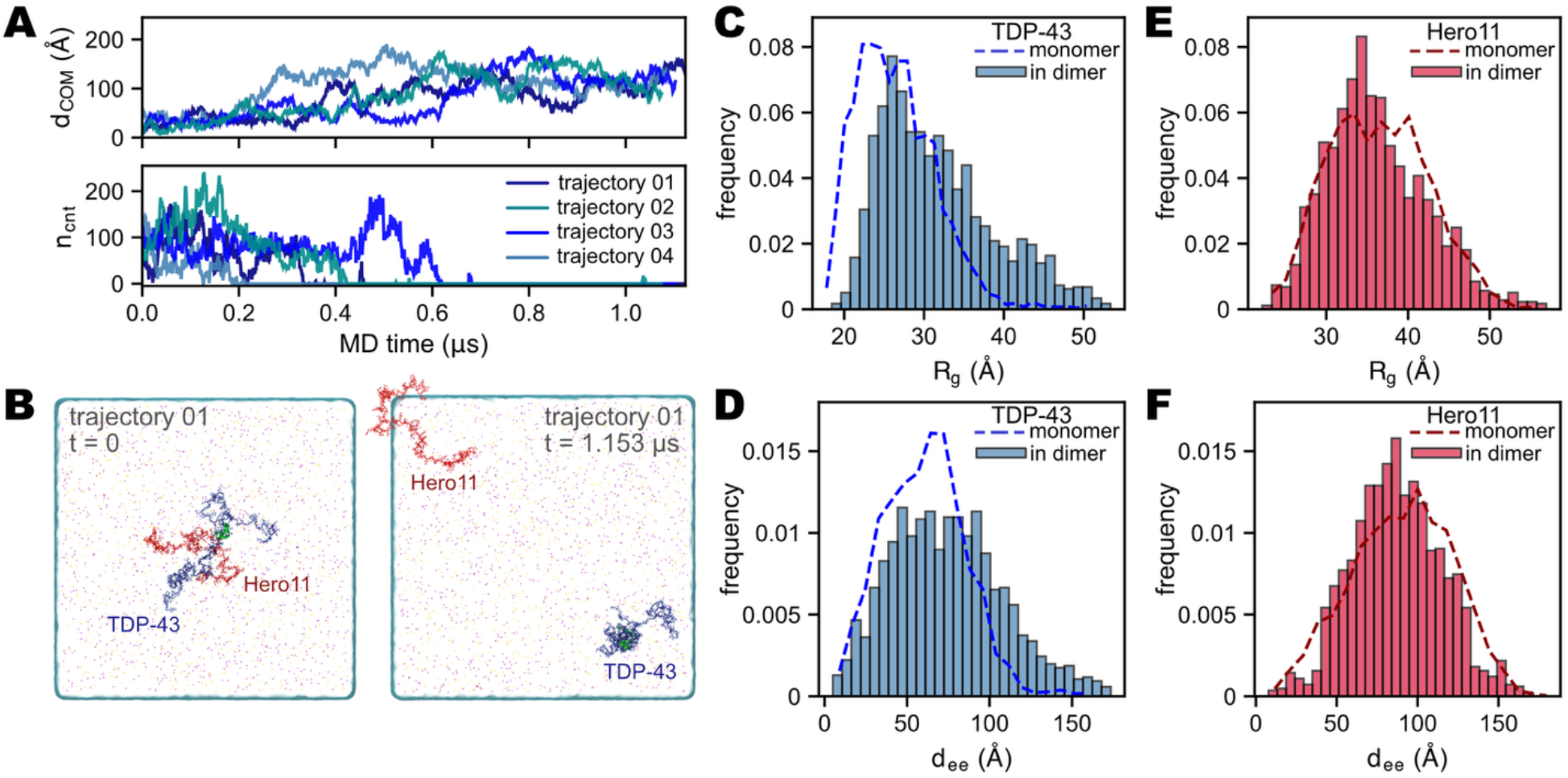
Simulations of TDP-43–Hero11 heterodimers in dilute solution. (A) Time evolution of the center-of-mass distance (*d*_*COM*_, upper panel) and the number of intermolecular contacts (*n*_*cnt*_, lower panel, see Methods for definition) between TDP-43 and Hero11 across four independent trajectories. Most trajectories show progressive dissociation of the heterodimer, with increasing *d*_*COM*_ and decreasing *n*_*cnt*_ over the course of =~1 μs simulations. (B) Representative snapshots from trajectory 01 at the beginning (*t* = 0, left) and end (*t* = 1.153 μs, right) of the simulation, illustrating the dissociation of the TDP-43 (blue)–Hero11 (red) complex. Ions are shown as small dots. (C, D) Distributions of the radius of gyration (*R*_*g*_, panel C) and end-to-end distance (*d*_*ee*_, panel D) of TDP-43 in the heterodimer states (bars, only using data when *n*_*cnt*_ > 0) compared with those from monomer simulations (dashed lines; here, C is the same as the distribution in Figure 2A). TDP-43 adopts more extended conformations upon binding to Hero11. (E, F) Same as (C, D) but for Hero11. The conformational properties of Hero11 remain largely unchanged between the monomer and dimer states.

Although stable binding was not observed, the simulations provide information on the conformational properties of TDP-43 in the presence of Hero11. To isolate frames in which the two chains were in contact, we selected trajectory snapshots with at least one intermolecular contact (*n*_*cnt*_ > 0), hereafter referred to as the “bound” state, and compared the resulting radius of gyration (*R*_*g*_) and end-to-end distance (*d*_*ee*_) distributions against the isolated monomer reference (Figure 3C–F). TDP-43 in the bound state adopted more extended conformations than in the monomer simulations, with both the *R*_*g*_ and *d*_*ee*_ distributions shifted toward larger values (Figure 3C, D). In contrast, the conformational properties of Hero11 were largely insensitive to whether it was in contact with TDP-43 (Figure 3E, F). The extension of TDP-43 upon transient association with Hero11 is qualitatively consistent with the single-molecule FRET measurements, that Hero11 shifts the conformational distribution of TDP-43 LCD toward more extended states^12^.

### Density and composition of homotypic and heterotypic condensates

To quantify the effects of Hero11 on the condensate, we computed the density profiles of all molecular species along the z-axis in the homotypic and heterotypic systems and examined their differences (*D*_*hetero*_ – *D*_*homo*_; Figure 4C–F). In the condensate core region, the TDP-43 density was lower in the heterotypic system than in the homotypic system, while the water density was correspondingly higher (Figure 4E). Notably, the total density (including all molecular species) in the core was also lower despite the additional presence of Hero11, indicating that Hero11 incorporation leads to a net decrease in the overall packing density of the condensate. The time evolution of these density profiles confirmed that both systems maintained stable condensate structures throughout the =~3.5 μs simulations (Figure S3).

**Figure 4.**
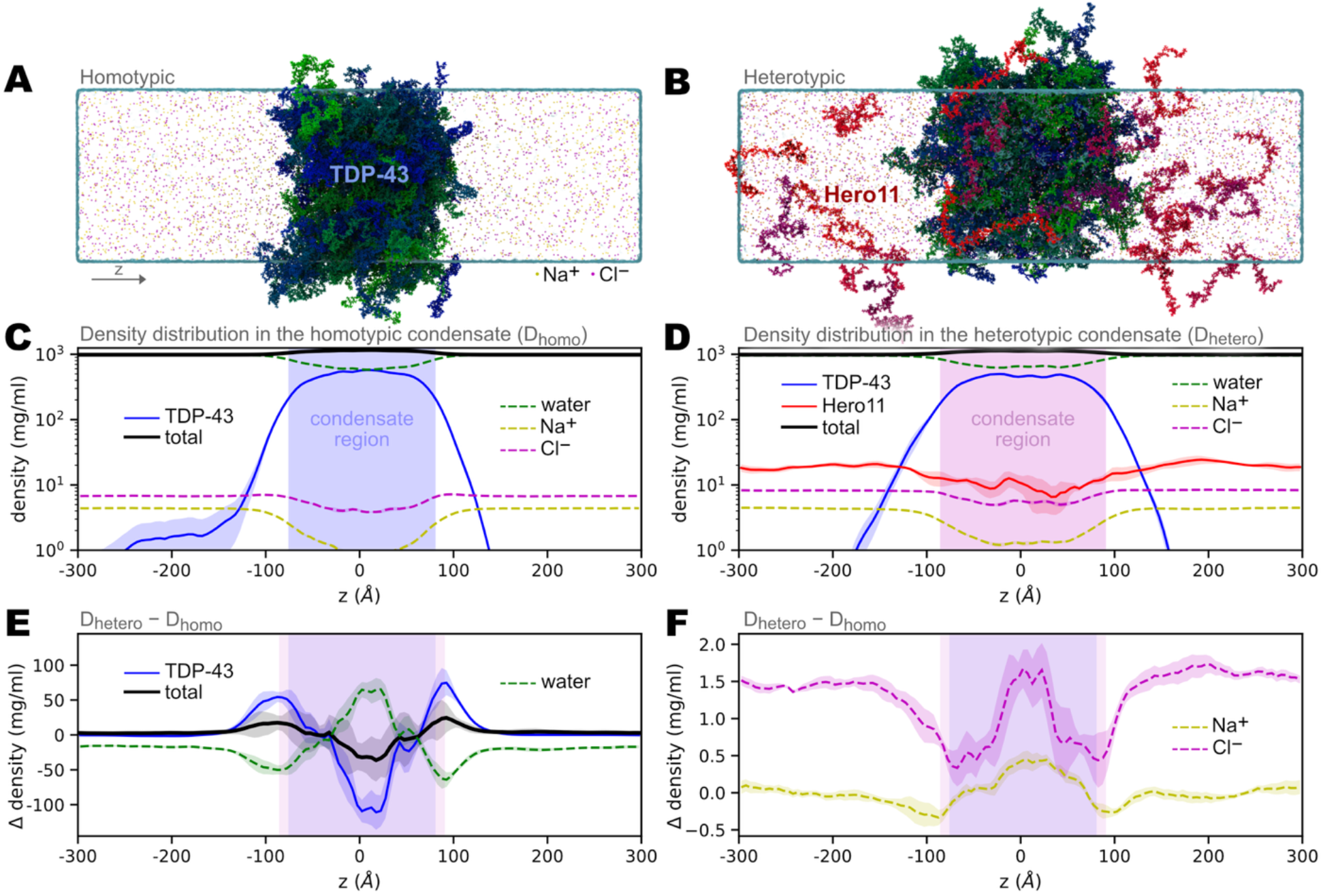
Density distributions in the homotypic TDP-43 and heterotypic TDP-43–Hero11 condensate systems from all-atom MD simulations. (A, B) Representative snapshots of the homotypic TDP-43 condensate (A) and the heterotypic TDP-43–Hero11 condensate (B). TDP-43 chains are shown in blue/green, Hero11 chains in red/crimson, Na^+^ ions in yellow, and Cl^−^ ions in green dots. The z-axis arrow indicates the longest dimension of the simulation box along which the density profiles were computed. (C) Density profiles of TDP-43 (blue solid line), water (green dashed line), Na^+^ (yellow dashed line), Cl^−^ (magenta dashed line), and total (black solid line) along the z-axis in the homotypic system (*D*_*homo*_). The shaded region indicates the condensate core (see Methods for definition). (D) Same as (C) but for the heterotypic system (*D*_*hetero*_), with the addition of the Hero11 density profile (red solid line). (E) Difference in density profiles (*D*_*hetero*_ – *D*_*homo*_) for TDP-43, total protein, and water. A decrease in TDP-43 density and an increase in water density are observed in the condensate region upon incorporation of Hero11. (F) Same difference plot as (E) but for Na^+^ and Cl^−^. Both ion species show increased concentrations within the condensate region in the heterotypic system. Shaded bands in (C–F) represent standard deviations calculated across independent trajectories.

Both Na^+^ and Cl^−^ showed elevated concentrations within the condensate core in the heterotypic system compared to the homotypic system (Figure 4F), despite their opposite charges. A more detailed analysis of the ion–residue interactions is presented in a later section.

### Hero11 disrupts the helix–helix interaction network

To identify the structural basis for the changes observed in the condensate density profiles, we analyzed the interchain contact probability between TDP-43 chains. The absolute contact probability maps for the homotypic and heterotypic systems are shown in Figure S4A and S4B, respectively. In both cases, the highest contact probabilities were concentrated around the α-helical region (residues 60–71), confirming that the helix serves as a major hub for interchain interactions within the condensate. The difference map (*ΔP* = *P*_*hetero*_ – *P*_*homo*_; Figure 5A) revealed that the most pronounced reduction in interchain contact probability occurred at the intersection of the helical regions, indicating that helix–helix contacts between TDP-43 chains are preferentially diminished in the presence of Hero11.

**Figure 5.**
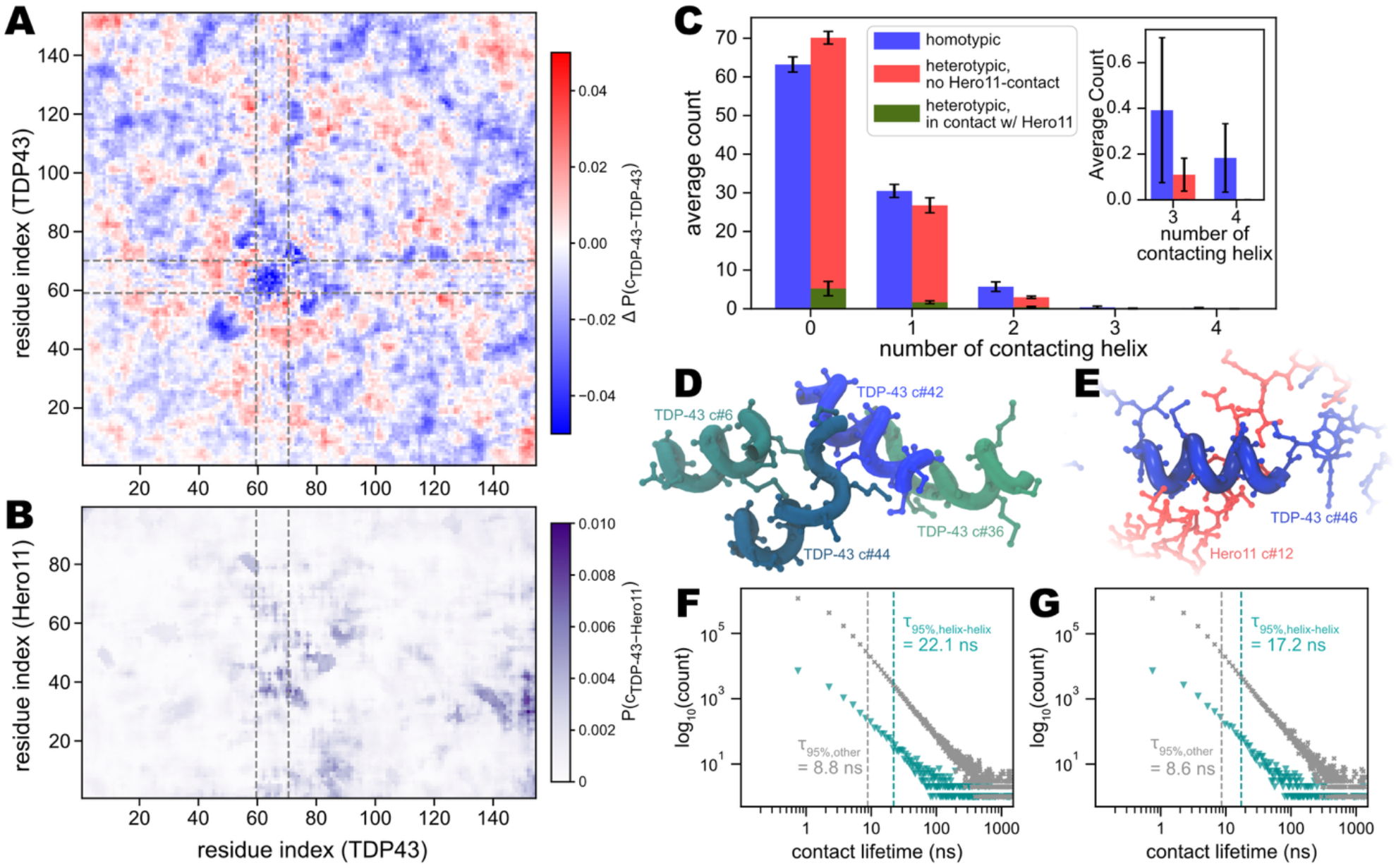
Hero11 disrupts the helix–helix interaction network in the TDP-43 condensate. (A) Difference in interchain contact probability between TDP-43 chains in the heterotypic versus homotypic condensates, *ΔP*(*c*_*TDP*−43−*TDP*−43_) = *P*_*hetero*_ – *P*_*homo*_. Blue indicates decreased contact probability upon Hero11 incorporation. Dashed lines indicate the α-helical region (residues 60–71). A pronounced reduction in contact probability is observed at the helix–helix intersection. (B) Interchain contact probability between TDP-43 and Hero11, *P*(*c*_*TDP*−43−*Hero*11_), in the heterotypic condensate. Dashed lines indicate the α-helical region of TDP-43. Hero11 exhibits preferential binding near the helical region as well as the C-terminal region of TDP-43. (C) Distribution of the helix–helix coordination number (number of other TDP-43 α-helices in contact with a given helix) in the homotypic condensate (blue), and in the heterotypic condensate separated into TDP-43 chains not in contact with Hero11 (red) and those in contact with Hero11 (dark green). Inset: magnified view for coordination numbers 3 and 4. The presence of Hero11, particularly in direct contact, reduces the helix–helix coordination number. (D) Representative snapshot from the homotypic condensate showing four TDP-43 chains (c#6, c#42, c#36, c#44, where c#*N* denotes the *N*-th chain) forming a helix–bundle arrangement. α-Helical regions are rendered as ribbons. (E) Representative snapshot from the heterotypic condensate showing Hero11 c#12 (red/pink) bound near the α-helical region of TDP-43 c#46 (blue). (F) Distribution of interchain contact lifetimes in the homotypic condensate. Helix–helix contacts (cyan triangles) exhibit longer lifetimes than non-helix contacts (gray crosses). Dashed vertical lines indicate the 95th percentile of each distribution (*τ*_95%_), with *τ*_95%,*helix*−*helix*_= 22.1 ns and *τ*_95%,*other*_ = 8.8 ns. (G) Same as (F) but for the heterotypic condensate. The helix–helix contact lifetime is reduced (*τ*_95%,*helix*−*helix*_ = 17.2 ns) compared to the homotypic case, while non-helix contact lifetimes remain similar (*τ*_95%,*other*_ = 8.6 ns). Error bars in (C) represent standard deviations across independent trajectories.

To understand where Hero11 binds within the condensate, we computed the interchain contact probability between TDP-43 and Hero11 residues (Figure 5B). The contact probability was the highest near the α-helical region of TDP-43, indicating that Hero11 preferentially interacts with this region. The binding was not restricted to a specific segment of Hero11 but was distributed across its sequence.

We further quantified the helix–helix interaction network by computing the coordination number, defined as the number of other TDP-43 α-helices in contact with a given helix (Figure 5C). In the homotypic condensate, a substantial fraction of helices had coordination numbers of 1 or higher, with some reaching 3 or 4 (Figure 5C, inset). In the heterotypic system, we separately analyzed TDP-43 chains that were in contact with Hero11 and those that were not. These results showed an increase in the fraction of helices with coordination number 0 and a corresponding decrease at higher coordination numbers compared to the homotypic case.

In the homotypic condensate, multiple (as many as four) TDP-43 helices could form closely packed arrangements (Figure 5D), whereas in the heterotypic system, Hero11 was found in the vicinity of the TDP-43 helical region, with fewer helix–helix contacts in the neighbourhood (Figure 5E). The packing geometry of helix–helix pairs was further characterized by the radial distribution function (RDF) and the joint distribution of distance and orientation between helix axes (Figure S5B–D). The RDF showed structured peaks at short distances in both systems, but with reduced amplitude in the heterotypic condensate (Figure S5B). The joint distance-orientation distributions indicated that closely spaced helix pairs favored parallel or antiparallel orientations in both systems, but perpendicular configurations were less populated in the heterotypic case (Figure S5C, D).

Finally, we examined the kinetics of interchain contacts by computing contact-lifetime distributions (Figure 5F, G). In the homotypic condensate, helix–helix contacts exhibited substantially longer lifetimes than non-helix contacts, with the 95th percentile of the lifetime distribution (*τ*_95%_) at 22.1 ns versus 8.8 ns (Figure 5F). In the heterotypic condensate, the helix–helix contact lifetime was reduced (*τ*_95%,*h*−*h*_ = 17.2 ns), while the non-helix contact lifetime remained similar (*τ*_95%,*other*_ = 8.6 ns) (Figure 5G). This selective reduction in helix–helix contact lifetime is consistent with the disruption of the helix–helix interaction network by Hero11.

### Changes in ion contacts, helical propensity, and condensate dynamics

We next examined how Hero11 incorporation affects the ion environment, secondary structure, and dynamics of TDP-43 within the condensate.

The per-residue difference in ion–residue contact probability (*ΔP* = *P*_*hetero*_ – *P*_*homo*_) is shown in Figure 6A. Both Na^+^ (upper panel) and Cl^−^ (lower panel) exhibited increased contact probabilities with TDP-43 residues in the heterotypic condensate. While the elevated Cl^−^ contacts were concentrated near positively charged residues (Arg/Lys), the increase in Na^+^ contacts was observed not only near negatively charged residues (Asp/Glu) but also at other positions along the sequence (Figure 6A). The latter observation is consistent with previous findings that Na^+^ can coordinate with backbone carbonyls in IDP condensates^20^. These results are also consistent with the increased overall ion concentrations within the condensate core observed in the density analysis (Figure 4F).

**Figure 6.**
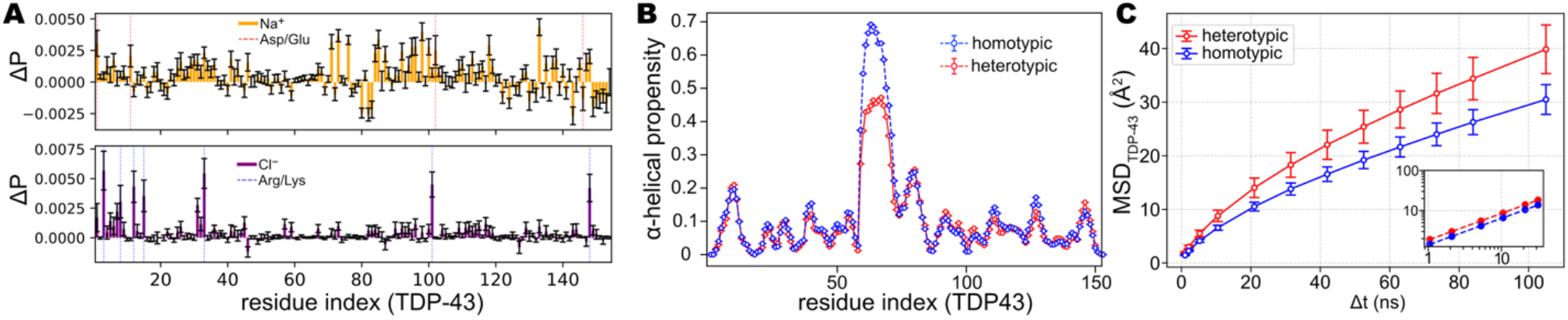
Effects of Hero11 on the ion distribution, secondary structure, and dynamics of TDP-43 in the condensate. (A) Difference in ion–residue contact probability (*ΔP* = *P*_*hetero*_ – *P*_*homo*_) mapped along the TDP-43 sequence. Upper panel: Na^+^ (orange bars) and positions of negatively charged residues (Asp/Glu, yellow dashed lines). Lower panel: Cl^−^ (purple bars) and positions of positively charged residues (Arg/Lys, blue dashed lines). (B) Per-residue α-helical propensity of TDP-43 in the homotypic (blue) and heterotypic (red) condensates. (C) Mean squared displacement (MSD) of the center of mass of TDP-43 chains in the dense phase as a function of time lag (*Δt*) for the homotypic (blue) and heterotypic (red) condensates. TDP-43 diffuses faster in the heterotypic condensate. Inset: the same data on a log-log scale. At short time lags (*Δt* < 40 ns), MSD follows a subdiffusive power law (MSD =~ *AΔt*^α^) with similar exponents (*α* ≈ 0.66) in both systems, but a larger prefactor in the heterotypic case. At longer time lags (*Δt* > 40 ns), MSD becomes approximately linear in *Δt*, with a larger slope in the heterotypic condensate (0.28 ± 0.02 Å^2^/ns) than in the homotypic condensate (0.22 ± 0.01 Å^2^/ns). Dashed lines in the inset represent the corresponding power-law fits. Error bars represent standard deviations across independent trajectories.

We then assessed the secondary structure of TDP-43 within the condensate by computing the per-residue α-helical propensity (Figure 6B). In the homotypic condensate, the region around residues 60–71 exhibited the highest helical propensity, reaching =~0.65. In the heterotypic condensate, the α-helical propensity of this region was reduced. This suggests that the crowded condensate environment stabilizes the helical structure (compared with the single-chain conformation, Figure S2D) but that this stabilization is partially diminished when Hero11 is present.

Finally, we computed the mean squared displacement (MSD) of the center of mass of TDP-43 chains in the dense phase (Figure 6C). At short time lags (*Δt* < 40 ns), the MSD followed a subdiffusive power law (MSD =~ *AΔt*^α^) with similar exponents in both systems (*α* ≈ 0.66), but with a larger prefactor in the heterotypic case. At longer time lags (*Δt* > 40 ns), the MSD became approximately linear in *Δt*, with a larger slope in the heterotypic condensate (0.28 ± 0.02 Å^2^/ns) than in the homotypic condensate (0.22 ± 0.01 Å^2^/ns). Thus, TDP-43 diffuses faster in the heterotypic condensate across all observed timescales.

## Discussion

### Weak pairwise interactions and a “kiss-and-run” model for Hero11 function in dilute solution

Our dimer simulations revealed that the pairwise interaction between TDP-43 and Hero11 is weak, with the heterodimer dissociating within the timescale of =~1 μs in most trajectories (Figure 3). Nevertheless, during the transient binding period, TDP-43 adopted more extended conformations compared to its monomeric state, which is qualitatively consistent with the smFRET measurements, which showed that Hero11 shifts the conformational distribution of TDP-43 LCD toward low-FRET, extended states^12^. Notably, the smFRET experiments were performed with Hero11 present as a soluble additive at 0.3 mg/mL, which corresponds to a large molar excess over individual TDP-43 molecules. In the original study, the aggregation-suppressing and neuroprotective effects of Hero proteins were also observed under overexpression conditions^11^. These observations, combined with the weak binding seen in our simulations, suggest a “kiss-and-run” model: individual Hero11 chains bind TDP-43 only transiently, but a sufficiently high concentration of Hero11 in the surrounding solution ensures that TDP-43 is frequently visited by Hero11, thereby maintaining the extended conformation on a time-averaged basis. This model implies that the protective effect of Hero11 in dilute solution is concentration-dependent and relies on the kinetic availability of Hero11, rather than on thermodynamically stable complex formation.

### Hero11 as a modulator that alters condensate properties through multivalency

A major finding of this study is that, despite the weak pairwise interaction observed in dilute solution, Hero11 remains associated with TDP-43 within the condensate (Figure 4B, D; Figure 5B). This indicates that the multivalent environment of the condensed phase is essential for sustaining effective Hero11–TDP-43 interactions. In this context, Hero11 functions as a condensate-incorporated modulator that enters the scaffold condensate and alters its physical properties from within^21,22^.

Ghosh et al. proposed three archetypical classes of macromolecular regulators of condensate phase behavior: volume-exclusion promoters, weak-attraction suppressors, and strong-attraction promoters^23^. In their framework, weak-attraction suppressors partition into the condensate but displace the stronger scaffold– scaffold interactions with weaker ligand–scaffold interactions, resulting in net destabilization. Hero11 shares some features with this class, in that it binds TDP-43 through relatively weak interactions and suppresses condensate density. However, Hero11 also carries a high net positive charge, and our previous CG simulations showed that electrostatic self-repulsion among Hero11 chains is an important driving force for loosening the condensate. We therefore suggest that Hero11 may represent an additional regulatory mode that could be termed a “repulsive suppressor”: a ligand that partitions into the condensate via weak ligand–scaffold attractions but destabilizes the condensate primarily through repulsive ligand–ligand interactions and disruption of the scaffold interaction network, rather than through simple competitive displacement. It is worth noting that the regulatory effect of Hero11 is itself multivalent: it simultaneously reduces helix–helix coordination, decreases helical propensity, and facilitates ion and water infiltration, effectively dismantling the same type of multivalent interactions that hold the condensate together.

The overall mechanism by which Hero11 regulates TPD-43 condensation as a repulsive suppressor, as inferred from our simulation, is summarized in Figure 7. In dilute solution, TDP-43 adopts a compact conformation with a transiently formed α-helix, while Hero11 remains extended due to electrostatic self-repulsion (Figure 7A). In the homotypic condensate, the α-helix is stabilized by the crowded environment and forms a percolated helix–helix interaction network that serves as the structural backbone of the dense phase (Figure 7B). Upon incorporation of Hero11, this network is partially disrupted through preferential binding of Hero11 near the helical region, accompanied by reduced helix–helix coordination, decreased helical propensity, increased water and ion infiltration, and enhanced protein diffusion (Figure 7C). We note that these changes co-occur as correlated observations in our simulations; whether they reflect a sequential causal cascade, a common upstream driver, or a combination of both remains an open question.

**Figure 7.**
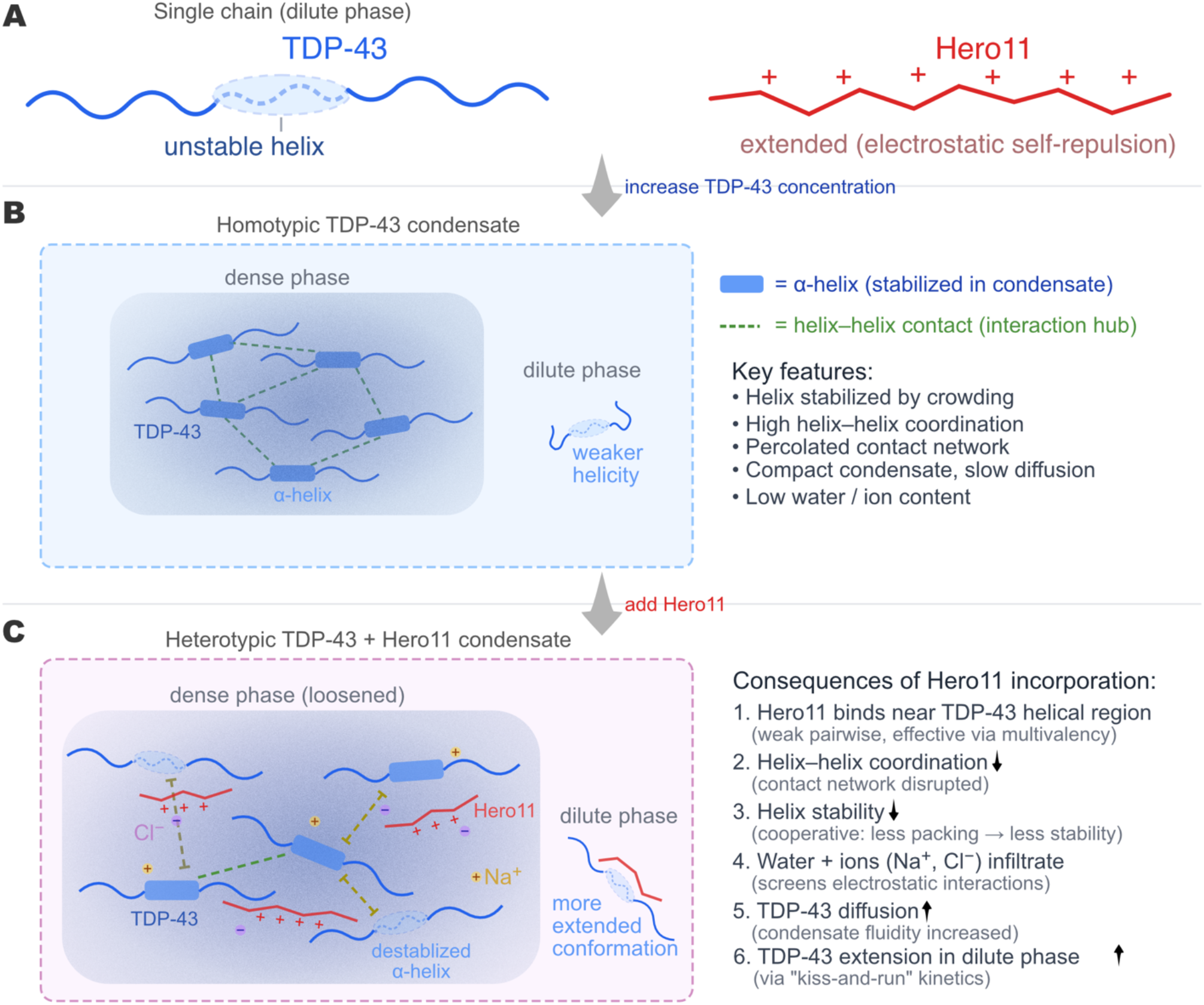
Mechanism for Hero11-mediated disruption of TDP-43 condensates. (A) In dilute solution, TDP-43 (blue) adopts a compact conformation with a transiently formed α-helix (dashed ellipse), while Hero11 (red) maintains an extended conformation due to intramolecular electrostatic self-repulsion among its abundant positive charges. (B) In the homotypic TDP-43 condensate, the α-helical region (blue rectangles) is stabilized by the crowded environment and forms a percolated helix–helix interaction network (green dashed lines) that serves as structural hubs. The resulting condensate is compact, exhibits slow protein diffusion, and has low water and ion content. (C) Upon incorporation of Hero11, the dense phase becomes loosened, accompanied by a series of correlated changes: (1) Hero11 preferentially binds near the α-helical region of TDP-43 through weak pairwise interactions that become effective via multivalency in the condensed phase; (2) the helix–helix coordination number is reduced, and the interaction network is partially disrupted; (3) the α-helical propensity of TDP-43 is concomitantly decreased; (4) water molecules and ions (Na^+^, Cl^−^) show increased concentrations within the condensate; (5) TDP-43 diffusion is enhanced, consistent with a more fluid condensate.

Our findings may also offer a partial answer to the question of how Hero proteins achieve client specificity. Experimental evidence indicates that different Hero proteins protect distinct client proteins, and no single Hero protein appears to be universally protective^11^. A purely electrostatic, non-specific mechanism, in which a highly charged protein simply repels aggregation-prone chains, would not fully account for this selectivity. Our simulations show that Hero11 does not interact uniformly with TDP-43 but instead preferentially associates near the transiently helical region (Figure 5), engaging it through a combination of electrostatic and aromatic contacts. This suggests that semi-structured elements within client proteins, rather than sequence-nonspecific charge complementarity alone, may contribute to target recognition by Hero proteins. We speculate that different Hero proteins may exploit analogous mechanisms, targeting distinct structural motifs in their respective clients. This hypothesis is testable and warrants systematic comparison across the Hero family.

### Simultaneous enrichment of Na^+^ and Cl^−^ in the heterotypic condensate

The observation that both Na^+^ and Cl^−^ concentrations increase in the heterotypic condensate (Figure 4F) is not straightforwardly explained by simple charge-based arguments. If the effect were purely electrostatic, one would expect the two ion species to respond in opposite directions upon incorporation of the highly positively charged Hero11. Instead, the simultaneous increase of both ion species is more consistent with an overall loosening of the condensate, which allows greater solvent and ion access to the interior. At the per-residue level, the enhanced Na^+^ contacts were observed not only near negatively charged residues (Asp/Glu) but also at other positions along the sequence (Figure 6A). This is consistent with the findings of MacAinsh et al. in their all-atom simulations of A1-LCD condensates, in which Na^+^ was found to coordinate preferentially with backbone carbonyls in addition to charged side chains, suggesting that cation–protein interactions in disordered protein condensates extend beyond simple electrostatic pairing^20^. In our system, the increased ion concentrations within the heterotypic condensate may further screen electrostatic interactions among TDP-43 chains, contributing to the observed reduction in condensation.

### Condensate dynamics hierarchy and comparison with electrostatically driven coacervates

In electrostatically driven coacervates of oppositely charged IDPs, inter-residue contacts are uniformly short-lived, with median lifetimes below 1 ns, and not rate-limiting for chain reconfiguration^24,25^. The TDP-43 condensate presents a qualitatively different picture. The contact lifetime distribution is markedly heterogeneous: helix–helix contacts exhibit 95th-percentile lifetimes of =~22 ns in the homotypic condensate (Figure 5F), roughly an order of magnitude longer than the median value reported for ProTα–H1 coacervates^24^, whereas non-helical contacts (=~8.8 ns) fall closer to that timescale. This hierarchy indicates that the α-helical network functions as a set of comparatively long-lived structural nodes within a background of more rapidly exchanging interactions. Furthermore, while oppositely charged IDPs cooperatively drive condensation through mutual electrostatic attraction^26,27^, Hero11 operates through the opposite mechanism: its high positive charge destabilizes the condensate. In the heterotypic system, helix– helix contact lifetimes decrease from =~22.1 ns to =~17.2 ns (Figure 5F, G), accompanied by increased translational diffusion (Figure 6C). The selective shortening of the longest-lived contacts, rather than a uniform acceleration, suggests a disproportionate effect on condensate stability from disrupting the slowest-exchanging elements of the network. These comparisons suggest that the relationship between contact lifetimes and condensate material properties established for polyelectrolyte coacervates extends to systems with more complex driving forces, but that the presence of structured interaction hubs, such as the α-helical network in TDP-43, introduces a distinct vulnerability to targeted disruption absent in uniformly charged systems.

### Comparison with coarse-grained simulations

The present all-atom simulations extend our previous CG study of TDP-43 and Hero11 condensates. At the CG level, we identified electrostatic repulsion between Hero11 chains as a key driver of destabilization of the condensate, and observed reduced interchain contacts and faster diffusion of TDP-43 in the presence of Hero11^15^. The all-atom simulations confirm these trends and, importantly, reveal molecular details that are inaccessible to CG models. First, the α-helical region of TDP-43 serves as a central structural hub within the condensate, forming a percolated helix–helix interaction network (Figure 5) that is stabilized by the crowded environment and absent under dilute conditions (Figure S2D). This finding could not have been anticipated from CG simulations, where secondary structure is either absent or imposed as a static input. Second, the all-atom simulations reveal that Hero11 preferentially targets the helical region (Figure 5B), providing a structural basis for its disruptive effect that goes beyond the electrostatic repulsion mechanism identified at the CG level. Third, the explicit treatment of water and ions reveals the simultaneous enrichment of both ion species within the heterotypic condensate (Figure 4F) and the ion–residue contact patterns at atomic resolution (Figure 6A), features that are inherently beyond the scope of implicit-solvent coarse-grained models.

## Conclusion

In this study, we performed large-scale all-atom MD simulations of TDP-43 condensates in the absence and presence of Hero11 to elucidate the molecular mechanism of Hero11-mediated condensate regulation. Our simulations revealed that the α-helical region of TDP-43 LCD, which is intrinsically unstable in the single-chain state, becomes stabilized within the condensate environment and forms a percolated helix–helix interaction network that serves as the structural hub of the condensate. Hero11 preferentially binds near this helical region, reducing the helix–helix coordination number and contact lifetime, thereby partially disrupting the interaction network. These changes are accompanied by a decrease in α-helical propensity, increased infiltration of water and both Na^+^ and Cl^−^ ions into the condensate core, and enhanced diffusion of TDP-43 within the dense phase. Dimer simulations further showed that the pairwise TDP-43–Hero11 interaction is weak under dilute conditions, suggesting that the regulatory effect of Hero11 operates through multivalent TDP-43–Hero11 interactions in the condensed phase rather than through strong binary binding. Together, these results identify the α-helix as a central structural element targeted by Hero11 and provide an atomic-level picture of how a repulsive ligand protein can modulate the physical properties of biomolecular condensates.

## Methods

### Construction of All-Atom Systems from Coarse-Grained Models

Initial all-atom (AA) configurations for TDP-43 and Hero11 were reconstructed from our previous CG MD simulations^15^ using a back-mapping procedure. The target proteins include the TDP-43 construct (containing a short α-helical segment and intrinsically disordered regions (IDRs)) and Hero11 (fully disordered). We selected representative snapshots from the CG trajectories of the condensate systems and extracted the coordinates of proteins located within or near the condensate interfaces. These selections typically included 100 TDP-43 chains and a varying number of Hero11 chains (17, 18, or 19, depending on the system stoichiometry).

We developed a hierarchical pipeline to convert Cα-only CG coordinates into full-atom models. First, individual protein chains were extracted from the CG snapshots. The all-atom structure of each chain was reconstructed using PULCHRA^18^, preserving the Cα positions from the CG model. Using these reconstructed chains, we built three categories of simulation systems: (i) Single-chain systems (3 independent replicates for TDP-43 and Hero11, respectively, Figure 1A); (ii) Dimer systems, where four pairs of TDP-43 and Hero11 in close proximity were identified from the dilute phase of the CG simulations (Figure 1B); and (iii) Condensate systems, where the reconstructed single chains were reassembled into the dense phase based on their original center-of-mass (COM) coordinates from the CG snapshots (Figure 1C).

To ensure structural integrity prior to solvation, we performed a geometric validation in vacuum using the structure-check utility in GENESIS^19^. This step resolved high-energy steric clashes, ring penetrations, and potential chirality errors introduced during the low-resolution-to-high-resolution mapping.

### Simulation Setup and Force Fields

All simulations were performed using the GENESIS software (version 2.1.0)^28–30^ on the supercomputer Fugaku. We employed the Amber99SBws-STQ force field for proteins^31^, which is optimized for intrinsically disordered proteins (IDPs) and protein-protein interactions, in conjunction with the TIP4P/2005 water model^32^ to accurately capture the hydration properties of disordered regions. The systems were solvated in cubic or rectangular boxes (see dimensions below) and neutralized with Na^+^ and Cl^−^ ions to reach a physiological salt concentration of 150 mM.

The simulation box dimensions were set as follows:

- Single-chain systems (Figure 1A): Cubic boxes with edge length of 220Å (TDP-43) and 180Å (Hero11).
- Dimer systems (Figure 1B): Cubic boxes with an edge length of 235 Å.
- Condensate systems (Figure 1C): Rectangular boxes of 180 × 180 × 600 Å for both homotypic and heterotypic condensates, with bulk water space to minimize finite-size effects perpendicular to the slab.

### Molecular Dynamics Protocol

The systems underwent a minimization and equilibration protocol to relax the backmapped structures and solvent environment. First, energy minimization was performed using the steepest descent algorithm, with water molecules treated as rigid. For single-chain and dimer systems, 5,000 minimization steps were used; for condensate systems, 20,000 steps were used to accommodate the larger system size and more complex initial geometry. The system was then heated from 0 K to 300 K over 50 ps in the canonical (NVT) ensemble. Subsequently, a 500-ps NVT equilibration was conducted at 300 K with positional restraints on protein heavy atoms (*k* = 2.0 kcal/mol/Å^2^).

For the isothermal-isobaric (NPT) equilibration^33^ at 300 K and 1 atm, we applied a stepwise relaxation of positional restraints to gradually adapt the system density. For single-chain and dimer systems, the restraints on protein heavy atoms were sequentially reduced (*k* = 2.0, 1.0, 0.5 kcal/mol/Å^2^) over three 0.5-ns stages, followed by a 5-ns restraint-free NPT run (total NPT duration: 6.5 ns). For condensate systems, a similar protocol was used but the 0.5 kcal/mol/Å^2^ restraint stage was extended to 2.5 ns, followed by a 5-ns restraint-free run (total NPT duration: 8.5 ns).

Production runs were performed in the NVT ensemble to maintain a constant simulation volume and a defined global macromolecular density. This setup is essential for accurately mapping phase behavior and prevents potential simulation-box instabilities arising from surface-tension effects at phase interfaces, which can occur in NPT simulations of heterogeneous slab systems.

We used the RESPA integration scheme^34^ (VRES in GENESIS). The time step was set to 3.5 fs, enabled by constraints on hydrogen-involved bonds (including SETTLE constraints for TIP4P water)^35–37^ and the Hydrogen Mass Repartitioning (HMR) with a scaling factor of 3.0 for solute hydrogen atoms^38^. Reciprocal-space electrostatic interactions were evaluated every 2 steps using particle mesh Ewald (PME) summation^39,40^. Real-space non-bonded interactions were calculated with a cutoff of 10.0 Å. Consistent with the force field parametrization, no switching function was applied to the van der Waals interactions. The temperature was maintained at 300 K using the stochastic velocity rescaling thermostat^41^ (*τ*_*T*_ = 5.0 ps), updated every 6 steps.

## Data Analysis

### Radius of gyration and conformational relaxation

The radius of gyration (*R*_*g*_) was computed using all non-hydrogen atoms. To estimate the characteristic timescale of conformational reconfiguration, we computed the normalized time autocorrelation function of *R*_*g*_:

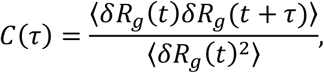

where *δR*_*g*_(*t*) = *R*_*g*_(*t*) − ⟨*R*_*g*_⟩ and angle brackets denote a time average over the production trajectory. *C*(*τ*) was then fitted to a single exponent decay,

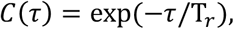

using nonlinear least-squares optimization, to obtain the characteristic relaxation time T_*r*_ (Figure 2H and 2J). The amplitude was fixed to unity to reflect the normalized autocorrelation.

#### Secondary structure assignment

Per-residue α-helical propensity was computed using the STRIDE algorithm^42^, as implemented in the ProteinSecondaryStructures.jl package. The α-helical propensity at each residue was defined as the fraction of trajectory frames assigned to an α-helical state.

#### Contact map and contact lifetime

Intramolecular contacts were defined by a Cα–Cα distance cutoff of 10 Å. This criterion was adopted in preference to a heavy-atom-based definition to reduce computational cost, which is particularly relevant for the multi-chain condensate systems analyzed using the same protocol (see below). Only non-local pairs were considered; pairs with sequence separation |*i* − *j*| < 4 were excluded.

For each residue pair (*i, j*), contact events were identified as contiguous intervals during which the Cα– Cα distance remained below the cutoff. The mean duration across all contact events for a given pair was taken as its representative lifetime (*τ*_*ij*_). The resulting per-pair mean lifetimes were then aggregated to construct the lifetime distribution. This per-pair averaging was chosen to weight each residue pair equally regardless of contact frequency, and to reflect a quantity analogous to the stability of a given interaction rather than the overall contact kinetics. To summarize the tail behavior of each lifetime distribution, we report the 95th percentile, *τ*_95%_, which characterizes the duration of the longest-lived contacts while remaining robust to outliers (Figure 2G, 2I, 5F, and 5G).

Pairs were classified into two categories: (i) helix–helix contacts, defined as pairs in which both residues *i* and *j* fall within the helical region (residues 60–71); and (ii) other contacts, comprising all the other pairs. Lifetime distributions for the two categories were analyzed separately for TDP-43 (Figure 2G). For Hero11 lifetime distribution does not distinguish the two categories.

For the intermolecular contacts in the condensate systems, we used the same Cα–Cα distance cutoff of 10 Å. Contact probability maps were computed for TDP-43–TDP-43 (Figure 5A) and TDP-43–Hero11 (Figure 5B) chain pairs, with each element representing the fraction of frames in which the corresponding residue pair was in contact, averaged over all chain pairs and independent trajectories.

Intermolecular contact lifetimes were computed following the same per-pair averaging protocol described above. For TDP-43–TDP-43 contacts, pairs were classified as helix–helix (both residues within indices 60–71) or other, consistent with the intramolecular analysis.

#### Helix-helix coordination

The helical region of TDP-43 was defined as residues 60–71 (corresponding to residues 320–331 in full-length TDP-43), based on the AlphaFold-predicted structure with per-residue pLDDT > 50. Note that this definition is sequence-based and was applied without reference to the instantaneous secondary structure assignment during the simulation, as the helix undergoes partial unfolding in some frames. For each chain *i*, the position of helix *i* was represented by the center of mass (COM) of all heavy atoms within the helical region, ***x***_*h,i*_, and its orientation by the unit vector 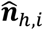 connecting the Cα atoms of residues 60 and 71. The pairwise helix–helix distance |***x***_*h,i*_, − ***x***_*h,j* |_ and orientational 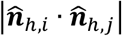 were computed for all pairs (*i, j*) with *i* ≠ *j* (Figure S5). The helix–helix coordination number of helix *i* was defined as the number of other helices *j* satisfying |***x***_*h,i*_ – ***x***_*h,j*|_ < 15 Å, a cutoff chosen to encompass the first shell of neighboring helices as indicated by the pairwise distance distribution (Figure S5B).

#### Density profile

To characterize the spatial distribution of each system component, the simulation box was divided into bins of width 5 Å along the z-axis (the longest box dimension). The local mass density of each component (TDP-43, Hero11, water, Na^+^, and Cl^−^) was computed in each bin for every trajectory frame. To account for translational diffusion of the condensate along z, each frame was aligned prior to averaging. Specifically, the total protein density profile was computed for each frame, and the center of the dense phase was identified as the midpoint of the region where the local TDP-43 concentration exceeded 3 mM. Each frame was then shifted along z such that this center coincided with the origin. The density profiles reported in Figure 4 represent averages over all frames and independent trajectories after this alignment. The boundaries of the condensate region were determined from the aligned, averaged total density profile. The ratio of successive bin densities, *ρ*_*i*_/*ρ*_*i*−1_, was computed along *z*, from −300 to 300 Å, and the positions of its maximum and minimum were taken as the two boundaries of the dense phase (shaded regions in Figures 4C–4F). For analyses requiring classification of individual protein chains as residing in the dense or dilute phase, each chain was assigned based on whether its center-of-mass *z*-coordinate fell within the condensate boundaries defined above.

#### Diffusion

The mean squared displacement (MSD) of TDP-43 was computed from the center-of-mass (COM) trajectories of TDP-43 chains residing in the dense phase. Chain position in the dense phase was evaluated at each frame; when a chain exited the condensate region, its trajectory was truncated at that point. The MSD was therefore computed from an ensemble of trajectory segments of varying lengths. For each replica, the MSD was averaged over all such segments, and the reported values represent the mean and standard deviation across the three independent replicas (Figure 6C).

## Acknowledgements

This work was supported in part by MEXT JSPS Kakenhi (grant number 19H05645, 21H05249 (to Y.S.), 21H05282 (to J.J. and C.T.), 25K09590 (to C.T.)), RIKEN pioneering projects “Biology of Intracellular Environments”, and “Glycolipidologue Initiative” (to Y.S.), RIKEN incentive (to J.J. and C.T.). MEXT program for promoting research on the supercomputer Fugaku (JPMXP1020200101), and MEXT program for Big-data-driven bio/synthetic polymer science to create absolutely circular materials (JPMXP1122714694) and Data-Driven Research Methods Development and Materials Innovation Led by Computational Materials Science (JPMXP1020230327) (to Y.S.). The computer resources are provided by the HPCI system research project (Project ID: ra000003, hp240090, HP260054) and by RIKEN Advanced Center for Computing and Communication (for HOKUSAI BigWaterfall, project RB230018).

## Notes

### Competing Interest Statement

The authors have declared no competing interest.

