## Supporting Figures for "Hero11 Unlocks TDP-43 Condensate Fluidity via Targeting Inter-Helical Interactions"

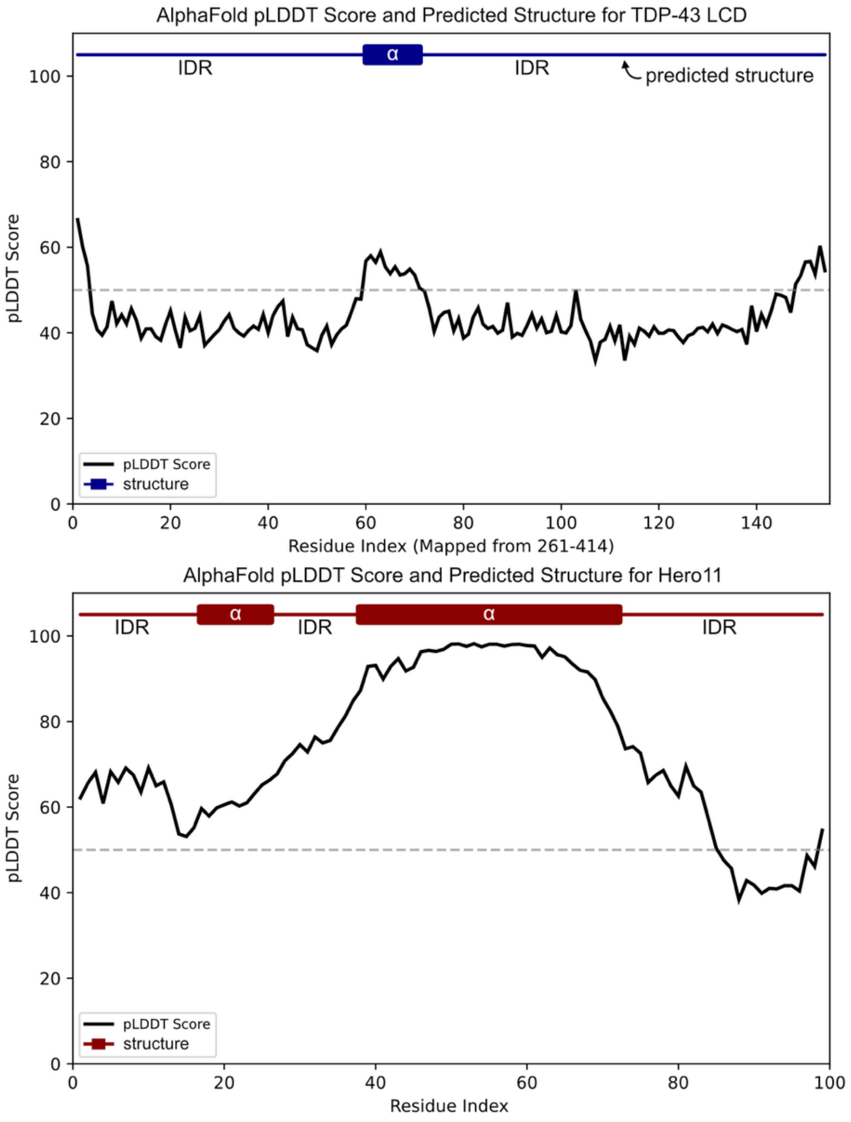


**Figure S1. AlphaFold-predicted structures and confidence scores for TDP-43 LCD and Hero11.** Upper panel: Per-residue pLDDT score and predicted secondary structure for TDP-43 LCD (residues 261–414, mapped to residue index 1–154). AlphaFold predicts a single α-helical region flanked by intrinsically disordered regions (IDRs), with moderately low confidence (pLDDT mostly below 50) throughout most of the sequence. Lower panel: Same analysis for Hero11. AlphaFold predicts two α-helical regions, with the longer helix (approximately residues 38–72) showing high confidence (pLDDT > 90) and the shorter helix (approximately residues 17–25) showing lower confidence (pLDDT ~ 60–70). The dashed horizontal line indicates pLDDT = 50.

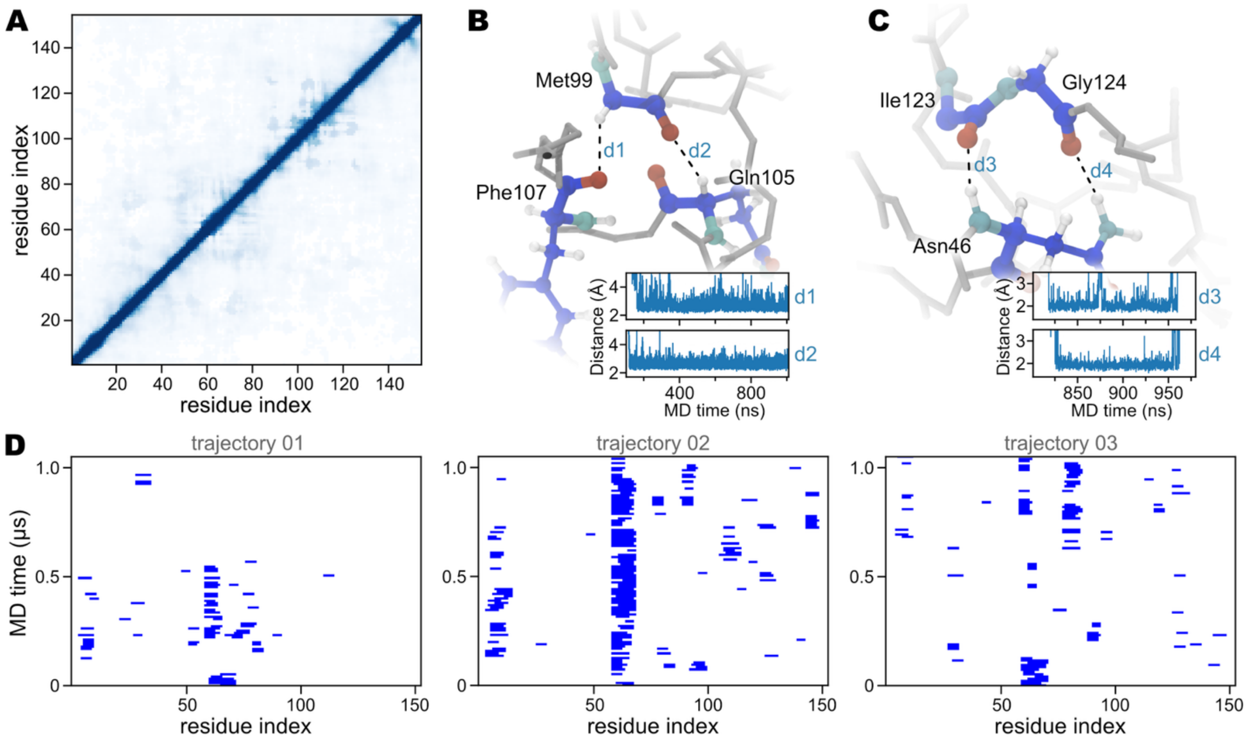


**Figure S2. Structural characterization of single-chain TDP-43 from all-atom MD simulations.** (A) Average intramolecular contact map of TDP-43. (B) Representative snapshot of a long-range intramolecular contact between Met99 and Phe107/Gln105. Dashed lines indicate monitored distances d1 and d2. Insets show the time evolution of d1 and d2 over the trajectory, demonstrating that this contact persists on the timescale of hundreds of nanoseconds. (C) Another example of a long-range intramolecular contact, between Ile123/Gly124 and Asn46. Insets show the time evolution of distances d3 and d4. The long-lived nature of such long-range contacts contributes to the compact conformation of TDP-43 in dilute solution. (D) Time evolution of per-residue α-helical assignment for three independent trajectories. Blue marks indicate that the residue is assigned as α-helix at the corresponding time point. Although the region around residues 60–71 exhibits the highest helical propensity, the α-helix undergoes repeated folding and unfolding events across all three trajectories, indicating that the helix is intrinsically unstable in the single-chain dilute condition.

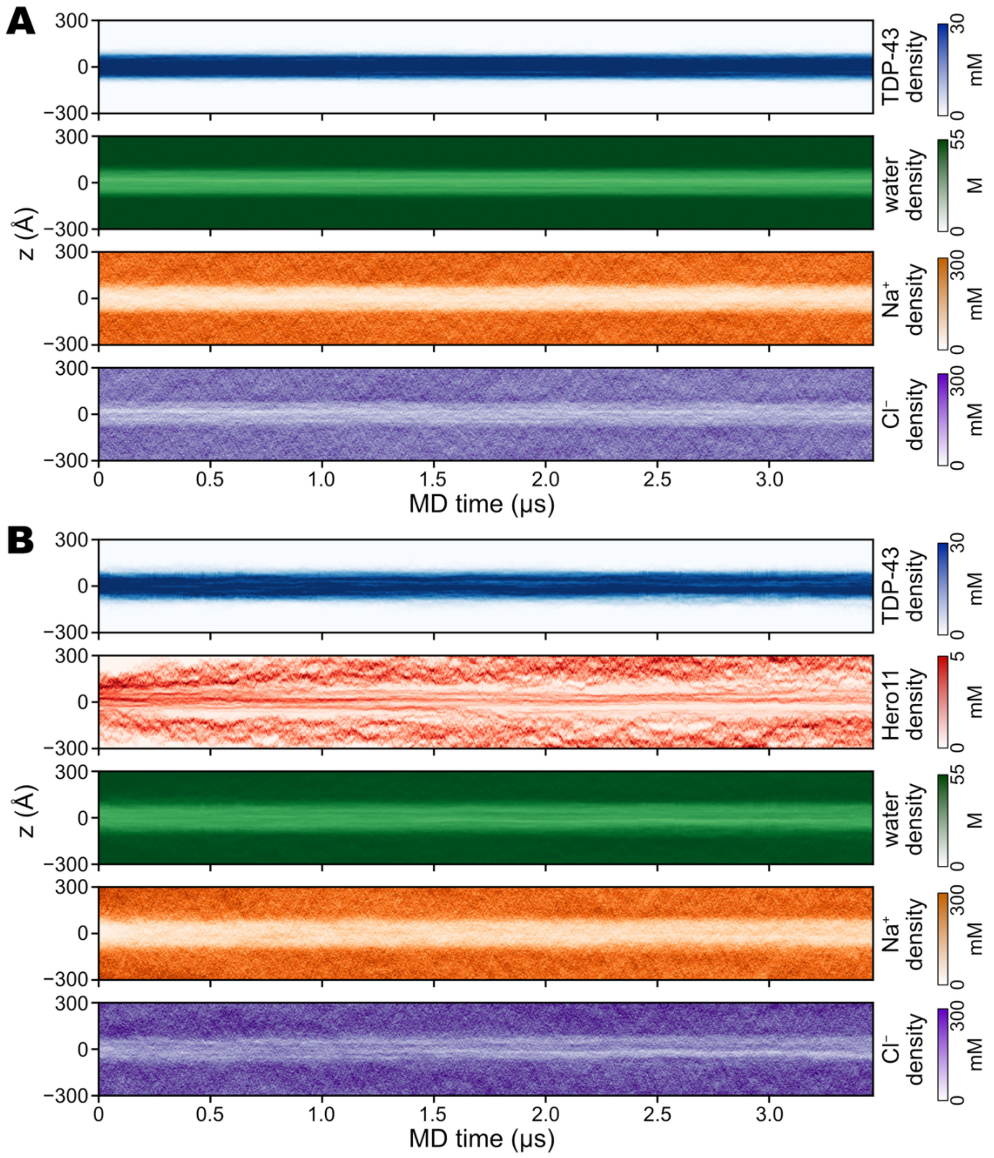


**Figure S3. Representative time evolution of density profiles along the z-axis in the condensate simulations.** (A) Homotypic TDP-43 condensate. From top to bottom: kymographs of TDP-43 density (blue, 0–30 mM), water density (green, 0–55 M), $Na^+$ density (orange, 0–300 mM), and $Cl^-$ density (purple, 0–300 mM) as a function of simulation time and position along the z-axis. (B) Heterotypic TDP-43–Hero11 condensate. From top to bottom: kymographs of TDP-43 density (blue, 0–30 mM), Hero11 density (red, 0–5 mM), water density (green, 0–55 M), $Na^+$ density (orange, 0–300 mM), and $Cl^-$ density (purple, 0–300 mM). Each panel shows the spatial distribution of the corresponding species over ~3.5 μs of simulation.

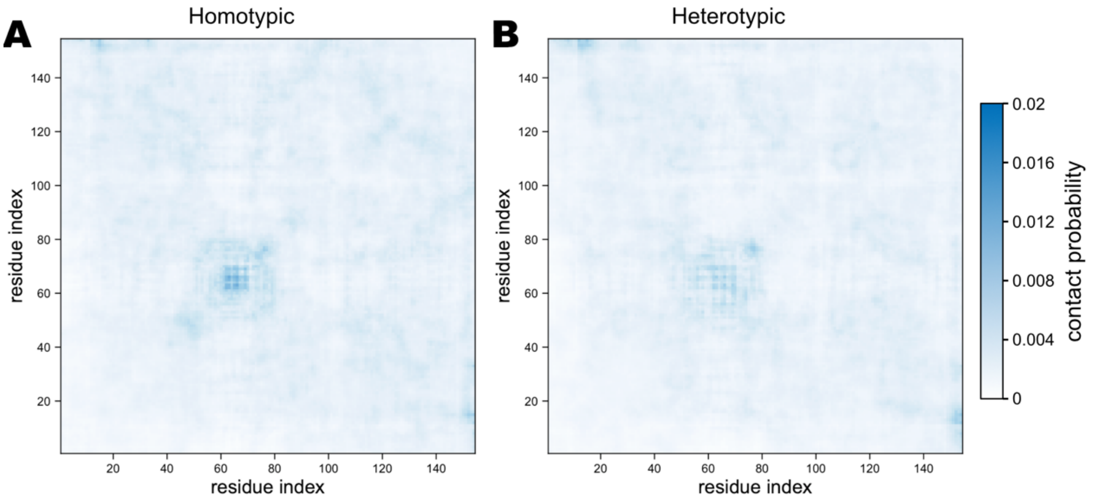


**Figure S4. Interchain contact probability maps between TDP-43 chains in the homotypic and heterotypic condensates.** (A) Interchain contact probability between TDP-43 residue pairs in the homotypic condensate. (B) Same as (A) but for TDP-43 chains in the heterotypic TDP-43–Hero11 condensate. Both maps share the same color scale. The highest contact probabilities are concentrated around the α-helical region (residues 59–69), appearing as a prominent feature near the diagonal block at this position. Figure 5A shows the difference map (B minus A).

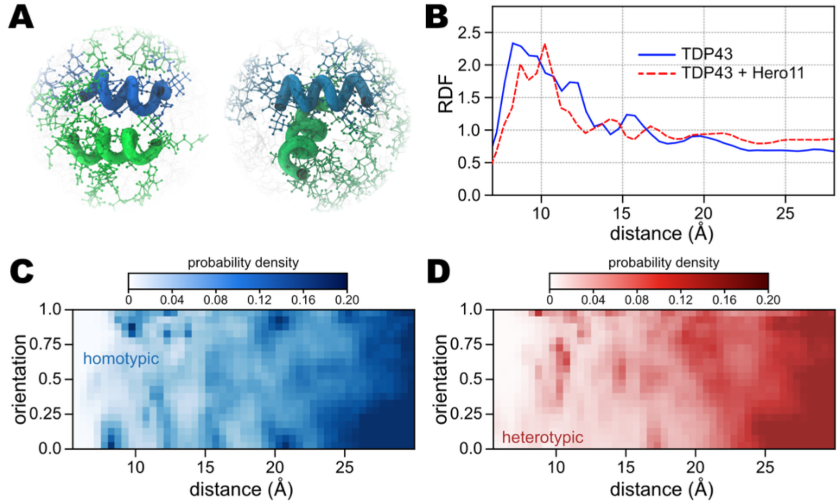


**Figure S5. Helix–helix packing geometry in the homotypic and heterotypic condensates.** (A) Representative snapshots illustrating the spatial arrangement of interchain helix–helix contacts between TDP-43 chains. Two examples are shown to illustrate typical packing geometries. (B) Radial distribution function (RDF) of the center-of-mass distance between α-helices from different TDP-43 chains in the homotypic condensate (blue solid line) and the heterotypic condensate (red dashed line). (C) Joint probability density of interchain helix–helix center-of-mass distance and orientation in the homotypic condensate. Orientation is defined as |cos θ|, where θ is the angle between the two helix axes, ranging from 0 (perpendicular) to 1 (parallel). (D) Same as (C) but for the heterotypic condensate.
